# Cyclic mechanical stretching stimuli promotes angiocrine signals during *in vitro* liver bud formation from human pluripotent stem cells

**DOI:** 10.1101/2023.06.11.544492

**Authors:** Koki Yoshimoto, Koichiro Maki, Taiji Adachi, Ken-ichiro Kamei

## Abstract

Liver organoids derived from human pluripotent stem cells (hPSCs) allow elucidation of liver development and have great potential for drug discovery. However, current methods for generating liver organoids using biochemical substances do not realize the vascular network of the liver lobule, due to the lack of knowledge of the role of *in vivo* mechanical environments during liver development. Here, we investigate the role of cyclic mechanical stretch (cMS) to angiocrine signals of hepatoblasts (HBs) and endothelial progenitor cells (EPCs) using an organ-on-a-chip platform to emulate *in vivo*-like mechanical environments and hPSCs to recapitulate hepatic differentiation. RNA sequencing revealed that the expression of angiocrine signal genes, such as *HGF* and matrix metallopeptidase 9 (*MMP9*), was increased by cMS in co-cultured HBs and EPCs. The secretion of HGF and MMP9 increased by 3.23-folds and 3.72-folds with cMS in the co-cultured HBs and EPCs but was not increased by cMS in the mono-cultured HBs and EPCs. Immunofluorescence micrographs with anti-KRT19, HGF, and MMP9 antibodies also revealed that cMS increased HGF and MMP9 expression when HBs and EPCs were co-cultured. cMS increased HGF and MMP9 expression and secretion when HBs and EPCs were co-cultured. Our findings provide new insights into the mechanical factors involved in the vascular network of human liver bud formation and liver organoid generation.

## Introduction

Liver development has attracted the attention of researchers and industries for a long time because of the fundamental interests in developmental biology and its applicability for disease modeling and drug discovery.^1–3^ In particular, human pluripotent stem cells (hPSCs), such as embryonic and induced pluripotent stem cells (hESCs^4^ and hiPSCs^5^ respectively), and their liver organoids allow the elucidation of liver development without the use of human embryos.^6,7^ Although the hepatic vascular network of the liver lobule is critical for expressing liver/hepatocyte functions, the vascular network in hPSC-derived liver organoids has not yet been established using current methods.^8–10^ During *in vivo* embryonic liver development, hepatoblasts (HBs) and endothelial progenitor cells (EPCs) differentiate from hepatic endoderm (HE) cells and migrate into the surrounding septum transversum mesenchyme to form a liver bud with the initial vascular network.^11,12^ Although the intercellular communication between HBs and EPCs plays an essential role in vascular network formation in a liver bud in mouse embryos.^13^ It is challenging to recapitulate the process of human hepatic vascular network formation *in vitro* due to a lack of fundamental understanding of human liver development.

In a previous study, human liver development using hPSC-derived organoids was conducted using biochemically soluble factors (growth factors and supplements) by recapitulating the extracellular environment of embryonic liver development.^11,14^ Because the foregut tube region, which produces the liver bud, is next to the fetal heart, liver development might also be influenced by the physical extracellular environment of the heart (cyclic stretching/pushing of the surrounding tissues and blood perfusion). Since blood perfusion does not occur during liver bud formation, mechanical stimuli of oscillated stretching/pushing by the heartbeat could be considered the main contributor to liver development. A recent study reported that mechanical stretching promotes the secretion of angiocrine signals to support the vasodilation and growth of human and mouse hepatocytes and endothelial cells *in vitro*.^15^ We hypothesized that cyclic mechanical stretching (cMS) promotes angiocrine signals in human adult liver regeneration and human liver bud formation via intercellular communication between HBs and EPCs.

To understand the influence of cMS on HBs and EPCs, establishing a highly controlled and recapitulated *in vitro* platform as an alternative model for a human embryo is necessary. The organ-on-a-chip (OoC) platform is advantageous over conventional *in vitro* cell-culture plates for generating and controlling cMS *in vitro* due to its ability to apply the mechanical forces and chemical gradients for cultured cells and organoids.^16,17^ We previously investigated the cMS generated by the OoC platform for HBs derived from hPSCs, which facilitated functionalities, such as cytochrome P450 3A (CYP3A),^18^ and demonstrated the applicability of the OoC platform for human liver development *in vitro*.

Here, we show that cMS activates angiocrine signals in hPSC-derived HBs and EPCs. Using the OoC platform, hPSCs were differentiated into HEs and EPCs, and cMS was applied to the cultured cells for three days. cMS for three days in a mixture of Hes and EPCs increased the expression of genes associated with epithelial-to-mesenchymal transition (EMT), including angiocrine signals in HBs and EPCs. We also investigated whether the activation of angiocrine signals requires intercellular interactions between HBs and EPCs via the secretion of EMT-associated proteins.

## Methods

### Microfluidic device fabrication

A microfluidic device was fabricated using stereolithographic 3D-printing techniques and solution-cast molding processes.^18, 19^ The molds for the top and bottom layers were produced using a 3D printer (Keyence Corporation, Osaka, Japan). After printing, the molds were washed with 99.9% ethanol for 12 h. The molds were then dried at 80°C for 30 min. A Sylgard 184 polydimethylsiloxane (PDMS) two-part elastomer (10:1 ratio of prepolymer to curing agent; Dow Corning Corporation, Midland, MI, USA) was mixed, poured into a 3D-printed mold to produce a 5-mm-thick PDMS top layer and a 2-mm-thick PDMS bottom layer, and degassed using a vacuum desiccator for 30 min. The PDMS was cured in an oven at 80°C for 16 h. After curing, the PDMS form was removed from the mold, trimmed, and cleaned. The Sylgard 184 PDMS two-part elastomer (10:1 ratio of prepolymer to curing agent) was poured onto a silicon wafer and spin-coated at 500 rpm for 30 s. After baking at 80°C for 10 min, the pressure chamber layer and PDMS thin membrane on the silicon wafer were treated with corona plasma (Kasuga Denki, Inc., Kawasaki, Japan) and subsequently placed at 80°C for 1.5 h for bonding. The bonded PDMS structure was then peeled off the silicon wafer. The top, bottom, and glass layers were corona-plasma-treated and bonded together by placing them in an oven at 80°C for 18 h. The devices were used within one day of completion.

### Device control

The PDMS membranes were actuated by airflow from a compressed air source (regulated at 0–200 kPa), operated with LabVIEW (version 11.0, National Instrument, Austin, TX, USA) software via solenoid valves (Takasago Electric, Inc., Osaka, Japan), using a controller board (VC3 8 controller [ALA Scientific Instruments, NY, USA] and NI USB-6501 [National Instruments, TX, USA]).

### hESC culture

The hESCs were used in accordance with the guidelines of the Ethics Committee of Kyoto University (ES3-9). H9 hESCs (WA09; RRID: CVCL_9773) were purchased from WiCell Research Institute (Madison, WI, USA). Before culturing, hESC-certified Matrigel (Corning, Inc., Corning, NY, USA) was diluted with DMEM/F12 (Merck KgaA, Darmstadt, Germany) at a 1:75 (v/v) ratio and coated onto a culture dish. Matrigel was incubated in a dish for 24 h at 4°C. Excess Matrigel was removed, and the coated dish was washed with fresh DMEM/F12. mTeSR-1-defined medium (Stem Cell Technologies, Vancouver, Canada) supplemented with 1% (v/v) penicillin/streptomycin (Fujifilm Wako, Osaka, Japan) was used for daily culturing of hESCs. For passaging, the cells were dissociated with TrypLE Express (Thermo Fisher Scientific, Waltham, MA, USA) for 3 min at 37°C and harvested. The cells were centrifuged at 200 ×*g* for 3 min, resuspended in the mTeSR-1 medium, and counted. mTeSR-1 medium containing 10 µM of the ROCK inhibitor Y-27632 (Fujifilm Wako) was used to prevent the apoptosis of dissociated hESCs on day 1. The mTeSR-1 medium without the ROCK inhibitor was used on subsequent days, with daily medium changes.

### Simultaneous endothelial and hepatic differentiation from hESCs on the device

Before inducing differentiation, the culture chambers of the device were coated with Matrigel at 35°C for 60 min. Matrigel was removed using an aspirator.

To induce endoderm differentiation, we washed cultured hESCs with D-PBS (no calcium, no magnesium) (Thermo Fisher Scientific) and treated them with TrypLE Express at 37°C for 3 min, followed by the addition of basal medium and transfer of the cell suspension into a 15-mL tube. The cells were centrifuged at 200 × *g* for 3 min, and the supernatant was removed. The cells were resuspended to 7.00 × 10^5^ cells mL^-1^ in mTeSR-1 medium supplemented with 10 µM Y27632, 100 ng mL^− 1^ activin A (human recombinant) (Fujifilm Wako), and 1% (v/v) penicillin/streptomycin. They were then applied to 30 µL chambers^-1^ resuspended solution in Matrigel-coated culture chambers and cultured in a humidified incubator at 37°C with 5% CO_2_ for 24 h. At the end of day 1, the medium was replaced with fresh mTeSR-1 medium supplemented with 10 µM Y27632, 100 ng mL^− 1^ activin A, and 1% (v/v) penicillin/streptomycin and cultured for an additional 24 h. On day 2, the medium was replaced with mTeSR-1 medium supplemented with 10 µM Y27632, 100 ng mL^− 1^ activin A, 10 ng mL^− 1^ BMP-4 (human recombinant) (R&D Systems, Minneapolis, MN, USA), 10 µM LY294002 (Cayman Chemical, Arbor, MI, USA), 3 µM CHIR99021(ReproCELL, Kanagawa, Japan), and 1% (v/v) penicillin/streptomycin. The cells were then incubated for 24 h. On day 3, the medium was replaced with mTeSR-1 medium supplemented with 10 µM Y27632, 100 ng mL^− 1^ activin A, 10 ng mL^− 1^ BMP-4, 10 µM LY294002, and 1% (v/v) penicillin/streptomycin, and the cells were incubated for 24 h. On day 4, the medium was replaced with Roswell Park Memorial Institute (RPMI) 1640 medium, GlutaMax Supplement (Thermo Fisher Scientific), supplemented with 2% (v/v) B-27™ supplement (Thermo Fisher Scientific), 1% (v/v) MEM Non-essential Amino Acid Solution without L-glutamine, liquid, sterile-filtered Bioreagent suitable for cell-culture (NEAA) (Merck KgaA), 1% (v/v) penicillin/streptomycin, 10 µM Y27632, 100 ng mL^− 1^ activin A, and 100 ng mL^− 1^ bFGF (human recombinant) (Fujifilm Wako), and the cells were incubated for 24 h.

To induce HE and EPC specification, we treated the cells with RPMI medium GlutaMax Supplement containing 2% (v/v) B-27 supplement, 1% (v/v) NEAA, 1% (v/v) penicillin/streptomycin, 10 µM Y27632, and 50 ng mL^− 1^ activin A, with daily media changes for three days. On day 8, to induce HB specification, the cells were treated with RPMI medium GlutaMax™ Supplement containing 2% (v/v) B-27™ supplement, 1% (v/v) NEAA, 1% (v/v) penicillin/streptomycin, 25 mM HEPES (Fujifilm Wako), 10 µM Y27632, 20 ng mL^− 1^ BMP-4, and 10 ng mL^− 1^ FGF-10 (human recombinant) (R&D Systems). The cells were then treated with RPMI medium GlutaMax™ Supplement containing 2% (v/v) B-27™ supplement, 1% (v/v) NEAA, 1% (v/v) penicillin/streptomycin, 25 mM HEPES, 20 ng mL^− 1^ BMP-4, and 10 ng mL^− 1^ FGF-10, with daily media changes and cMS for three days.

### RNA sequencing (RNA-seq)

Total RNA was purified using the Rneasy Micro Kit (Qiagen, Hilden, Germany) and frozen. NEBNext® Ultra™II Directional RNA Library Prep Kit for Illumina® and NEBNext Poly(A) mRNA Magnetic Isolation Module were used as library prep kits. The total PCR cycles were six. For RNA-seq analysis, reads were aligned to a human reference genome (GRCh38.p13.genome.fa), and the number of reads mapped to each gene and transcript was counted using HISAT2 and the featureCounts package with GENCODE (gencode.v36.annotation.gtf).^20,21^ Read count data were analyzed, and differentially expressed genes (DEGs) were identified with a false discovery rate <0.1 and |log_2_ fold change| >1 and produced with iDEP.96.^22^ Gene ontology (GO) analysis of the DEGs was performed, and GO terms were produced using Metascape ^23^ and TRRUST.^24^

### Cell sorting with a fluorescence-activated cell sorter (FACS)

The cells were rinsed with PBS twice and harvested using 0.1 % trypsin /EDTA and 1 mg mL^− 1^ trypsin inhibitor before cell counting. For antibody staining, cells were diluted to a final concentration of 1 × 10^7^ cells mL^− 1^ in staining buffer (fetal bovine serum) (BD Pharmingen, Franklin Lakes, NJ, USA); 2.5 µL of fluorescence-labeled antibodies was added to 50 µL of cell suspension, and incubated on ice for 30 min (Table S3). As a negative control, specific isotype controls were used with the same primary antibody concentration (Table S5). After removing excess antibodies by centrifugation at 300 ×*g* for 3 min, the cells were washed with staining buffer, and cell suspensions were applied to a FACSAria II SORP (BD Biosciences, Franklin Lakes, NJ, USA) for cell sorting. Data were analyzed using the FlowJo software (v9; FlowJo, LLC, Ashland, OR, USA).

### Cell culture on the device after cell sorting

After HBs and EPCs were sorted with KDR antibody, the cells were resuspended with RPMI medium GlutaMax Supplement containing 2% (v/v) B-27 supplement, 1% (v/v) NEAA, 1% (v/v) penicillin/streptomycin, 25 mM HEPES, 10 µM Y27632, 20 ng mL^− 1^ BMP-4, and 10 ng mL^− 1^ FGF-10; 1.0 × 10^4^ cells/well were seeded onto a device coated with Matrigel as a monoculture, and 1.0 × 10^4^ cells per well were directly seeded on the device after harvest as a co-culture.

### Sprouting assay

Geltrex 100 µL (Thermo Fisher Scientific) was added to the dish and incubated at 37°C for 30 min. The cells in the 16 wells of our device were sorted with the KDR antibody and seeded on the gel. The cells were cultured using the EGM Endothelial Cell Growth Medium BulletKit (Lonza, Basel, Switzerland) and observed after 48-h incubation to investigate whether the cells formed a sprouting structure unique to ECs.

### Reverse transcription-quantitative PCR (RT-qPCR)

Total RNA was purified using an RNeasy Micro Kit (Qiagen), and 12 ng of total RNA was reverse-transcribed to generate cDNA using PrimeScript RT Master Mix (Perfect Real Time; TaKaRa Bio, Shiga, Japan) after cell sorting. A reaction mixture (25 μL) containing 100 pg cDNA, 0.2 μM PCR primers (Table S6), and 5 U of Taq DNA polymerase (TaKaRa Bio) was subjected to PCR using a thermal cycler (Applied Biosystems 7300 real-time PCR system; Applied Biosystems, Foster City, CA, USA). PCR was performed with 30 to 40 cycles (94°C for 30 s, 58°C for 30 s, and 72°C for 60 s).

### Immunocytochemistry

The cells were fixed with 4% paraformaldehyde in D-PBS (-) (Fujifilm Wako) for 20 min at 25°C and permeabilized with 0.1% (v/v) Triton X-100 in D-PBS for 5 min at 25°C. Subsequently, the cells were blocked in D-PBS (5% (v/v) normal goat serum blocking solution (Maravai Life Sciences, San Diego, CA, USA), 5% (v/v) normal donkey serum (Jackson ImmunoResearch, West Grove, PA, USA), 3% (v/v) albumin, essentially globulin-free (Merck KgaA), and 0.1% Tween-20 (Nacalai Tesque, Kyoto, Japan) at 4°C for 16 h and incubated at 4°C for 16 h with primary antibodies in blocking buffer as described in Table S5. The cells were then incubated at 37°C for 60 min with a secondary antibody, as described in Table S7, before a final incubation with DAPI (Fujifilm Wako) at 25°C for 30 min.

### Protein quantification

The cells on the device were rinsed with cold D-PBS, and 25 µL/well cold 1× RIPA buffer (Cell Signaling Technology, Danvers, MA, USA) in double distilled water was added. The cells were incubated on ice for 30 min, collected into tubes, sonicated at 4°C (US-1R cleaner; AS ONE, Osaka, Japan), and centrifuged at 4°C, 10,000 ×*g* for 20 min. The supernatant was collected into new tubes and stored at -20°C. Using a BCA protein kit (TaKaRa, Shiga, Japan), a working solution (BCA reagent A: B = 100:1) and 0.2 mg µL^-1^ BCA standard were mixed with the collected proteins in 100 µL 1× RIPA buffer in a 96-well plate (Matrix Microplate w/lids 96-well blk/clr, flat bottom, tissue culture, PS; Thermo Fisher Scientific). Proteins were incubated at 37°C for 60 min and measured at a wavelength of 562 nm using a Synergy HTX Microplate Reader.

### ELISA

The culture supernatants incubated for 24 h after adding the medium were collected from the two wells to evaluate each sample’s HGF and MMP9 secretion capacity. Human HGF Quantikine ELISA (R&D, DHG00B) and Human MMP-9 Sandwich ELISA Kit (Proteintech, KE00164) were used for cell culture supernatants according to the manufacturer’s instructions. The amounts of secreted HGF and MMP9 were normalized by the protein content in the two wells.

### Image acquisition

The sample containing cells was placed on the stage of a Nikon ECLIPSE Ti inverted fluorescence microscope equipped with a CFI plan fluor 10×/0.30 N.A. objective lens (Nikon, Tokyo, Japan), CCD camera (ORCA-R2; Hamamatsu Photonics, Hamamatsu City, Japan), mercury lamp (Intensilight; Nikon), XYZ automated stage (Ti-S-ER motorized stage with encoders; Nikon), and filter cubes for fluorescence channels (DAPI, GFP HYQ, TRITC; Nikon). Immunochemical analysis was conducted using CellProfiler (version 3.1.9; Broad Institute of Harvard and MIT, USA).^25^ Otsu’s method ^26^ was used to identify the nuclei and cells, and fluorescence signals from individual cells were quantified automatically, except for cells touching the borders of the images.

### Statistical analysis

One-way ANOVA followed by Tukey’s test and two-way ANOVA followed by Welch’s test were performed using *R* software (ver. 4.0.2). Two-tailed, unpaired Student’s *t*-test was conducted with Microsoft Excel.

## Results

### OoC platform to apply cMS during liver bud formation from hESCs

The OoC platform was fabricated using soft lithography combined with a 3D printer.^18,19^ This platform has mainly three layers of PDMS: a cell culture layer, a stretching layer, and a pressure chamber layer (Figure 1A). The cell culture layer has 32 chambers, and the bottom chamber is a PDMS membrane as a stretching layer. As previously described,^18^ this OoC platform allows the mechanical stretching of cultured cells on the membrane by applying air pressure to the pressure chambers under the cell culture chambers. The degree of mechanical stretching is regulated by the pressure drop from the air inlet to the outlet. Cell culture chambers were coated with Matrigel to allow the adhesion, growth, and differentiation of H9 hESCs; 3.00 × 10^3^ H9 hESCs mm^-2^ in a cell culture chamber were seeded and differentiated into HEs and EPCs, and cMS was applied from day 9 (D9) to D12 (Figure 1B).

**Figure 1.**
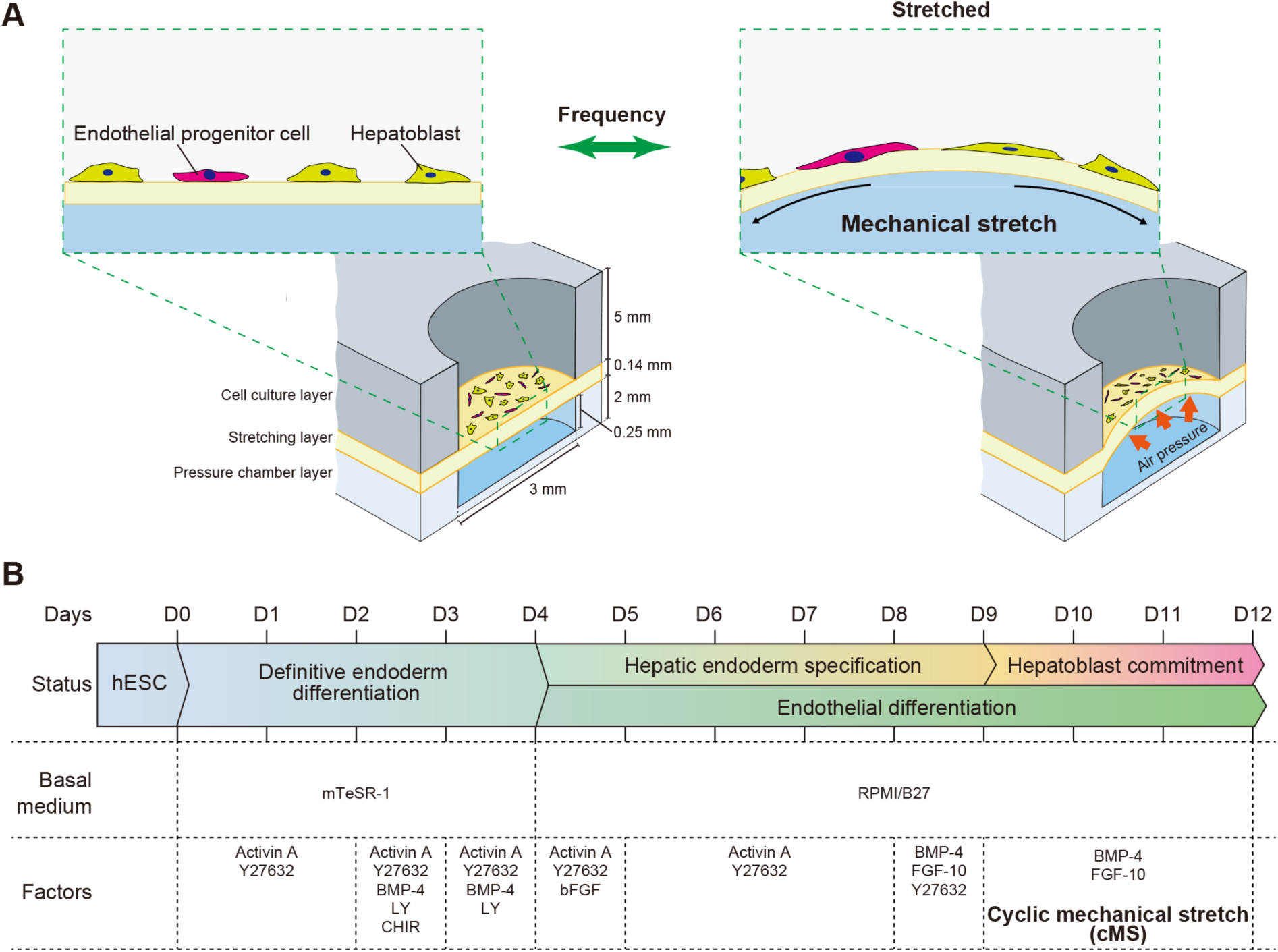
Overview of the contribution of cMS to HBs and EPCs using an OoC platform and hESCs. (A) Illustration of an OoC platform to apply cMS to HBs and EPCs derived from hESCs. (B) Schematic illustration of the simultaneous differentiation from hESCs to HBs and EPCs. cMS was applied after 9-day differentiation from hESCs. Abbreviations: cMS: cyclic mechanical stretching, EPC: endothelial progenitor cells, hESCs: human embryonic pluripotent stem cells, HB: hepatoblast, OoC: organ-on-a-chip

### HEs and EPCs were simultaneously generated from hESCs in the OoC platform

A recent report suggested that hESCs show mixed populations of HEs and EPCs after HE specification.^27^ We confirmed that the previously reported method for differentiating hESCs from HEs at D9^18, 28^ provided mixed populations of HEs and EPCs, even in the on-chip culture before cMS (Figures 2 and S1). The obtained HEs showed the expression of the HE markers, including keratin 19 (KRT19), forkhead box A2 (FOXA2), hematopoietically-expressed homeobox protein (HHEX), SRY-box transcription factor 17 (SOX17) and hepatocyte nuclear factor 4α (HNF4α), confirmed by immunocytochemistry (Figures 2A and S1A). In contrast, the obtained EPCs expressed a typical EC marker, KDR. Quantitative single-cell profiling based on the images in Figure 2A showed that HE and EC markers were exclusively expressed in HBs and EPCs, respectively (Figures 2B and S1B).

**Figure 2.**
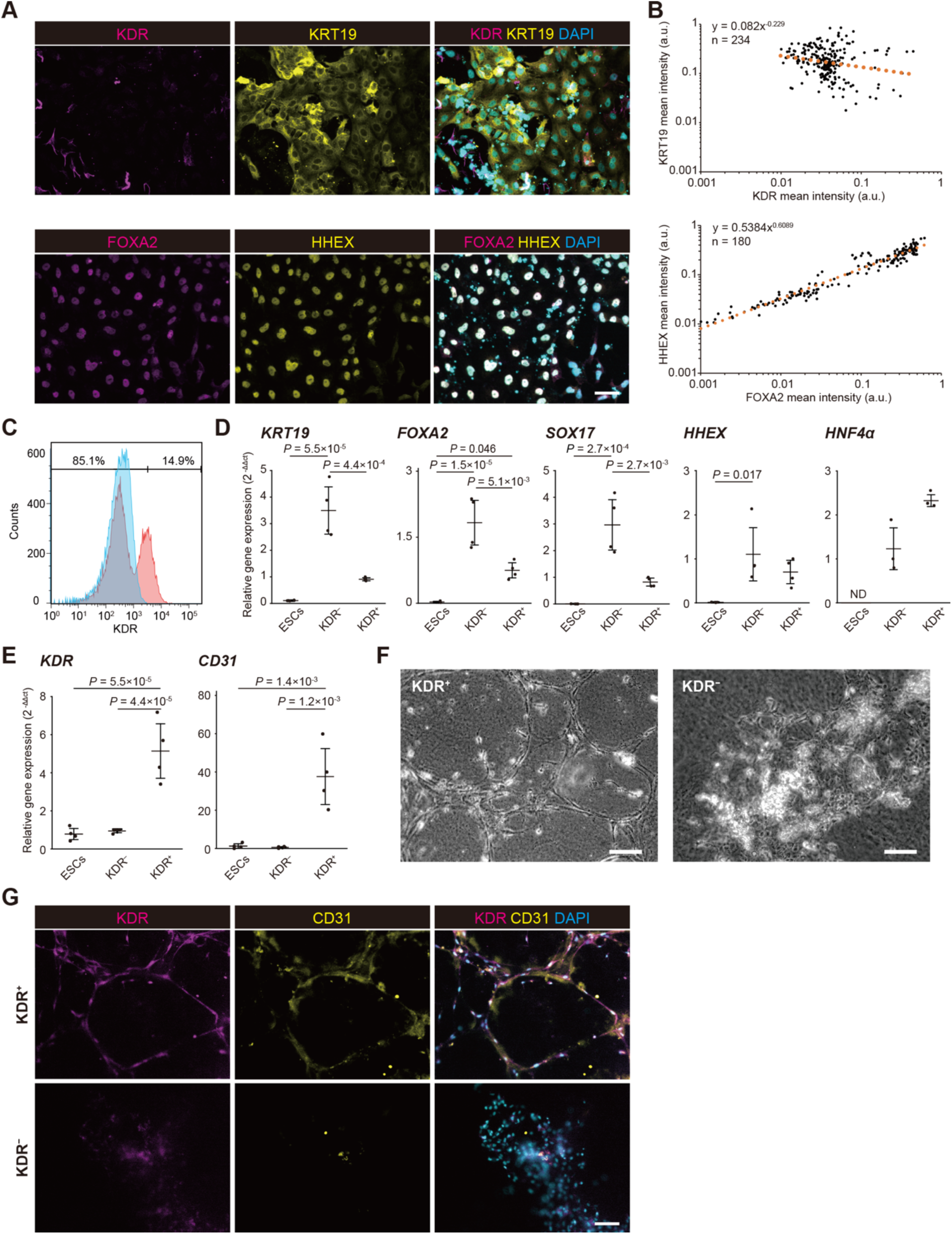
*In vitro* differentiation of H9 hESCs to HE and EPCs at D9. (A) Immunofluorescence micrographs of D9 cells of the mixture stained with the HE markers (KRT19, yellow; FOXA2, magenta; HHEX, yellow), the EC marker (KDR, magenta), and the nuclei (DAPI, cyan). Scale bars represent 100 µm. (B) Scatter plots showing the quantitative single-cell profiling of the expression of KDR/KRT19 (upper panel) and FOXA2/HHEX (bottom panel) based on fluorescent microscopic images shown in (A). The approximate lines were fitted to quantitative single-cell profiled results to show the correlative relationships. (C) Flow cytometric analysis showing KDR expression in the mixed populations of HE cells and EPCs derived from H9 hESCs on the OoC platform. (D, E) Real-time PCR showing the expression of HE (D, *KRT19, FOXA2, SOX17, HHEX*, and *HNF4α*) and EC (E, *KDR*, and *CD31*) markers in KDR^−^ and KDR^+^ cells after cell sorting. All data represent means ± SD. The significance value was calculated using one-way ANOVA followed by Tukey’s test. (n=4) (F) An endothelial sprouting assay of KDR^−^ and KDR^+^ cells on the gel-containing basement membrane after cell sorting. Scale bar represents 100 µm. (G) Immunofluorescence micrographs of KDR^−^ and KDR^+^ cells on the gel-containing basement membrane after cell sorting stained with the EC marker (KDR, magenta; CD31, yellow), and the nuclei (DAPI, cyan). Scale bars represent 200 µm. Abbreviations: D9: day 9, EPC: endothelial progenitor cells, FOXA2: forkhead box A2, HE:hepatic endoderm, hESCs: human embryonic pluripotent stem cells, HHEX: hematopoietically-expressed homeobox protein, HNF4α: hepatocyte nuclear factor 4α, KRT19: keratin 19, OoC: organ-on-a-chip, SOX17: SRY-box transcription factor 17

To further evaluate the expression of genes associated with HB and EC markers in D9 cells, we performed FACS using the KDR EC marker, followed by RT-qPCR (Figure 2C). For FACS analysis, D9 cells had 13.9 ± 1.42% of KDR-positive (KDR^+^) cells, while most KDR^+^ cells did not express the CD31 EC marker, suggesting that these cells were endothelial progenitor-like cells (Figure S1C and D). According to the RT-qPCR analysis, among the HE markers, *KRT19, FOXA2*, and *SOX17* levels were significantly lower in sorted KDR^+^ cells than in KDR-negative (KDR^−^) cells, although *HHEX* and *HNF4A* levels were not significantly different between KDR^−^ and KDR^+^ cells (Figure 2D). In contrast, KDR^+^ cells showed higher gene expression levels of *KDR* and *CD31* than KDR^−^ cells (Figure 2E). Furthermore, KDR^+^ cells showed an endothelial spouting formation in the gel-containing basement membrane, suggesting their functionality as ECs (Figure 2F).^29^ Interestingly, the KDR^+^ cells after sprouting formation expressed CD31 along the vascular networks, while KDR^−^ cells did not (Figure 2G). These results indicated that D9 KDR^−^ and KDR^+^ cells represented HEs and EPCs, respectively.

### RNA sequencing revealed cMS induced the expression of EMT-associated genes in hESC-derived HBs and EPCs

Previously, we reported that cMS with 0.1 Hz of frequency and a stretch at 575±42 µm of the membrane displacement in our OoC platform facilitated the CYP3A activities in HBs and EPCs derived from H9 hESCs at D12.^18^ Applying the same cMS from D9 to D12 during the differentiation of H9 hESCs, we conducted RNA-seq with or without cMS (D12_cMS^+^ and D12_cMS^−^, respectively) (Figures 3, S2-S4). RNA samples from H9 hESCs differentiated at D9 without cMS (D9_cMS^−^) were also analyzed for comparison. For the principal component analysis, PC1 distinguished the tested samples by differentiation stage at D9 and D12 (Figure 3A). In contrast, PC2 distinguished the D12 samples based on the effects of cMS.

**Figure 3.**
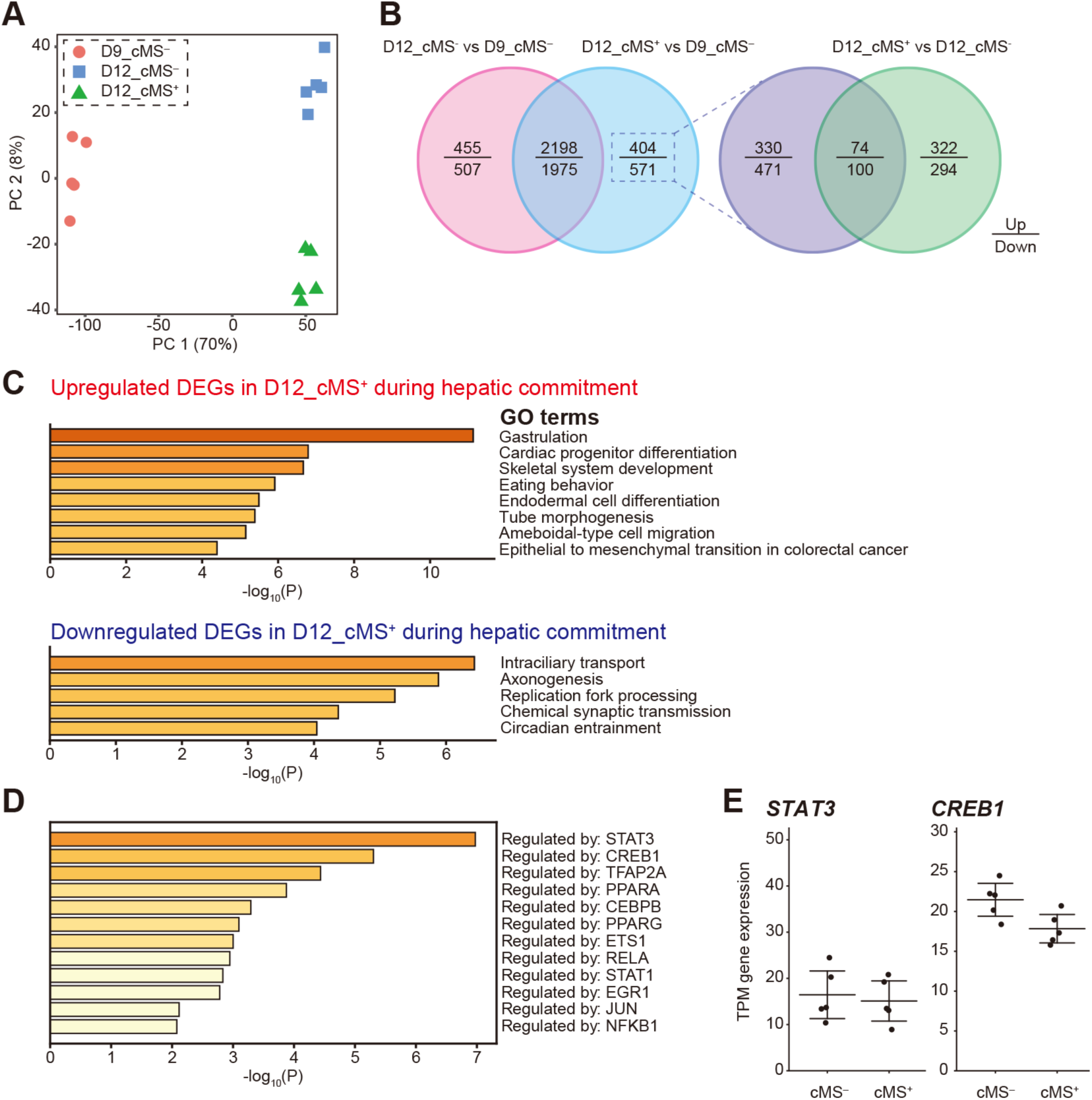
Comparative transcriptional profiling and secreted EMT-related proteins of D9, D12_cMS^−^ and D12_cMS^+^. (A) Principal component analysis based on the RNA-seq with D9, D12_cMS^−^, and D12_cMS^+^ showed that all the mixtures at D9 (pink), D12_cMS^−^ (blue), and D12_cMS^+^ (green) were segregated into three groups. (B) Venn diagrams comparing the identified DEGs in D12_cMS^−^/D9 and D12_cMS^+^/D9 (left) as well as the unique DEGs in D12_cMS^+^/D9 and D12_cMS^+^ vs. D12_cMS^−^. (C) Gene ontology terms based on 74 upregulated and 100 downregulated DEGs identified in (B). (D) Identification of transcription factors regulating 74 upregulated DEG. (E) The TPM gene expressions of *STAT3* and *CREB1* in D12_cMS^−^ and D12_cMS^+^. All data represent means ± SD. Abbreviations: cMS: cyclic mechanical stretching, D9: day 9, D12: day 12, DEG: differentially expressed gene, EMT: epithelial-to-mesenchymal transition, SD: standard deviation, TPM: transcripts per kilobase million

To investigate the role of cMS during HB commitment, we identified 4930, 4944, and 732 DEGs by analyzing D12_cMS^−^ vs. D9_cMS^−^ (Figure S2 and Table S1), D12_cMS^+^ vs. D9_cMS^−^ (Figure S3 and Table S2), and D12_cMS^+^ vs. D12_cMS^−^ (Figure S4 and Table S3), respectively. Based on the Venn diagram of DEGs in D12_cMS^−^/D9_cMS^−^ and D12_cMS^+^/D9_cMS^−^, 2198 upregulated and 1975 downregulated DEGs were shared between D12_cMS^−^/D9_cMS^−^ and D12_cMS^+^/D9_cMS^−^ ; however, 404 upregulated and 571 downregulated DEGs were observed only in D12_cMS^+^/D9_cMS^−^ (Figure 3B). Among the uniquely observed DEGs in D12_cMS^+^/D9_cMS^−^, 330 upregulated and 471 downregulated DEGs were observed in D12_cMS^+^/D9_cMS^−^ compared to 726 upregulated and 865 downregulated DEGs in D12_cMS^+^ vs. D12_cMS^−^, suggesting the cMS-dependent gene sets during HB commitment; 74 upregulated and 100 downregulated DEGs were observed as shared gene lists with D12_cMS^+^ vs. D12_cMS^−^, suggesting that they were cMS-dependent but were not involved in HB commitment (Table S4).

To obtain the biological insights of obtained DEGs of cMS-dependent gene sets during HB commitment, we explored their biological functions, found that upregulated DEGs have a collection of the EMT-related genes, including angiocrine-related genes (*NODAL, MMP9, MESP1, IL6, ZEB2*, and *TWIST1*)^30–35^ and also found the GO term associated with EMT (“Epithelial to mesenchymal transition in colorectal cancer”) (Figure 3C). TRRUST^24^ suggested that STAT3 and CREB1 were the main transcription factors for the upregulated DEGs (Figures 3D and S4B), whereas the expression of *STAT3* and *CREB1* was not changed by cMS (Figure 3E).

Moreover, cMS may activate continuous calcium ion flow into the cytoplasm through mechanosensitive channels such as PIEZO1 or transient receptor potential vanilloid 4 (TRPV4).^36,37^ Our transcripts per kilobase million (TPM) gene expression profiles showed higher gene expression levels for some PIEZO and TRP family members, such as *PIEZO1, TRPM4*, and *TPRM7* in the mixture (Figure S5). These mechanosensitive channels may be involved in the activation of SP1, STAT3, and CREB3. In contrast, we could not identify DEGs associated with cell-cell interactions (cadherins, connexins, and junctional adhesion molecules), which might also be altered by cMS. Thus, these analyses showed that cMS mainly upregulated EMT-related genes in HBs and EPCs, which were mainly regulated by STAT3 and CREB1 on D12.

### cMS enhanced the secretion of HGF and MMP from HBs and EPCs on D12

Among the EMT-related genes upregulated in D12_cMS^+^, *HGF* and *MMP9* play essential roles in hepatic progenitor cell migration and hepatic vascular network formation *in vivo*^38–40^; however, it is unclear how cMS regulates the production of HGF and MMP9 in HBs and EPCs. As *HGF* and *MMP9* expression is regulated by SP1 and STAT3, which might be activated through the mechanoreceptors sensitive to the frequency and magnitude of stretch, we measured the secretion of HGF and MMP9 from the cells with the frequency of 0.01, 0.1, and 1 Hz and the stretching at 447 ± 20, 505 ± 30, and 575 ± 42 µm after 24 h (Figures 4A and S6). The secretions were significantly increased corresponding to the increase of frequencies, and the 1 Hz frequency and stretch at 575 ± 42 µm displacement made HBs and EPCs at D12 secrete 3.8-fold higher HGF and 3.4-fold higher MMP9 than HBs and EPCs at D12 without cMS. Therefore, we used the 1 Hz frequency and a stretch of 575 ± 42 µm displacement for further analyses.

**Figure 4.**
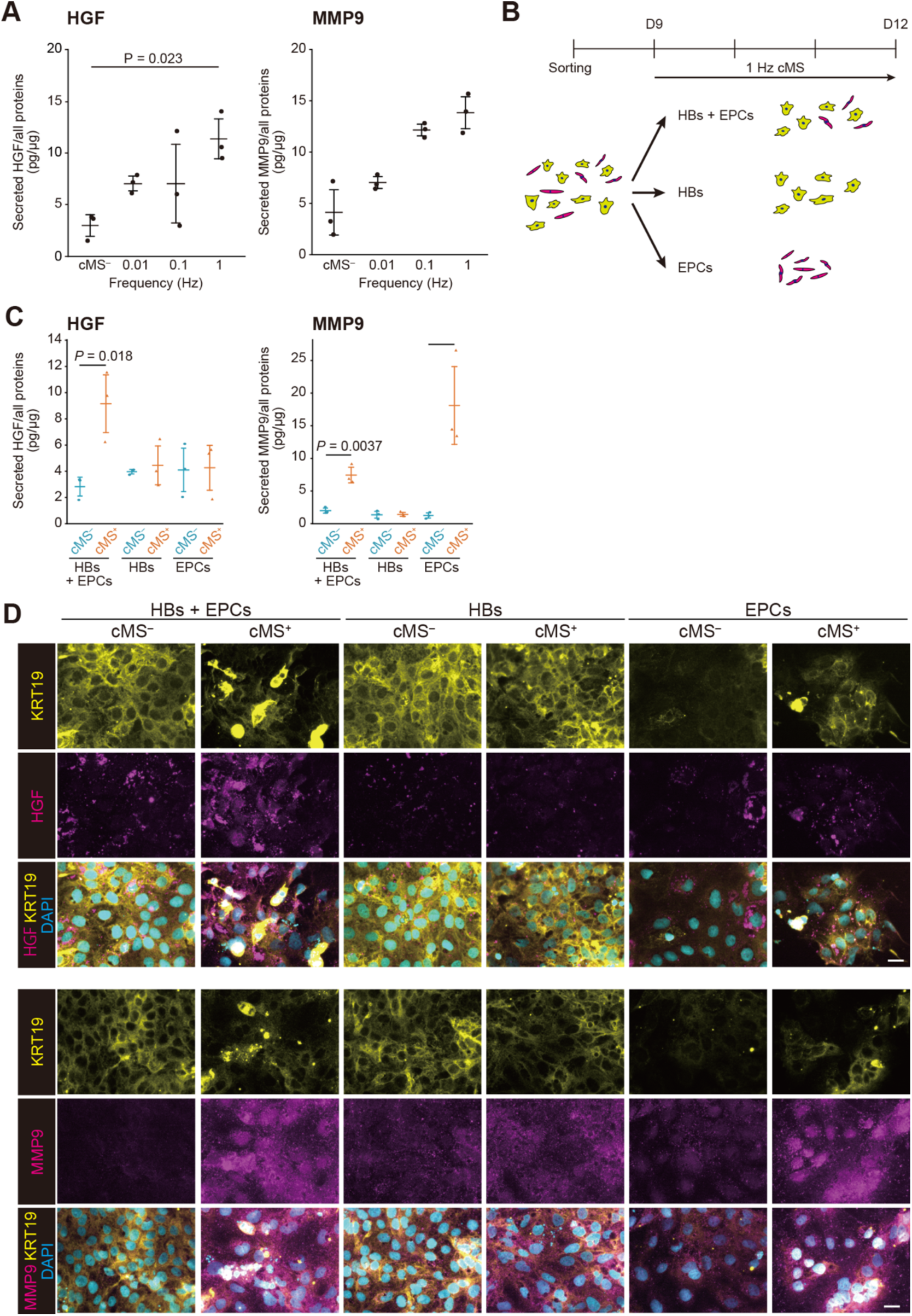
cMS enhanced the expression and secretion of EMT-associated HGF and MMP9 from HBs and EPCs. (A) Secretion of HGF and MMP9 in the mixtures of HBs and EPCs with 0.01, 0.1, and 1 Hz cMS and without cMS. All data represent means ± SD. The significance value was calculated using one-way ANOVA followed by Tukey’s test (n=3). (B) Schematic illustration to sort HBs and EPCs followed by cMS. (C) Secretion of HGF and MMP9 from the co-cultured and mono-cultured HBs and EPCs by applying cMS at D12. All data represent means ± SD. The significance value was calculated using a two-tailed unpaired Student’s *t*-test (n=4). (D) Immunofluorescence micrographs of HB marker (KRT19, yellow), EMT marker (HGF, magenta; MMP9, magenta), and the nuclei (DAPI, cyan) in co-cultured and mono-cultured HBs and EPCs at D12. Scale bars represent 25 µm. Abbreviations: cMS: cyclic mechanical stretching, D12: day 12, EMT: epithelial-to-mesenchymal transition, EPC: endothelial progenitor cells, HB: hepatoblast, MMP9: matrix metallopeptidase 9, SD: standard deviation

To investigate which HBs and EPCs at D12 secreted HGF and MMP9, we sorted hESC-differentiated cells at D9 by FACS with the KDR EC marker and applied cMS to the sorted cells (Figure 4B). We confirmed cMS did not alter the cellular populations of KDR^−^ and KDR^+^ cells at D12 (Figure S7A). According to RT-qPCR analysis, KDR^−^ cells expressed HB markers (*CYP3A7* and *HNF4A*), whereas KDR^+^ cells expressed the EC markers (*KDR* and *CD31*) (Figure S7B). Only <1% of KDR^+^ cells in D12_cMS^−^ and D12_cMS^+^ expressed CD31 EC markers, suggesting that these cells were endothelial progenitor-like cells (Figure S7C). Thus, KDR^−^ cells and KDR^+^ cells were termed HBs and EPCs, respectively.

The expressions of typical EMT-related genes (*HGF, MMP2, MMP9*, and *TWIST1*) were quantified in HBs and EPCs on D12 after cell sorting using the KDR EC marker (Figure S7D). By applying cMS, HBs increased *HGF* expression 43.7 folds, while EPCs did not show a significant difference with cMS. In particular, HBs and EPCs with cMS showed 22.0- and 12.0-fold increases in *MMP9* expression, respectively, compared to HBs and EPCs without stimulation.

HGF and MMPs are secreted proteins used for intercellular communication.^15^ To investigate whether HBs and EPCs required HGF and MMP9 to communicate with each other, we sorted HEs and EPCs independently seeded into the device. Although HGF secretion was increased 3.23-fold by cMS in co-cultured HBs and EPCs, HGF secretion was not changed by cMS in separately cultured HBs and EPCs (Figure 4C). In contrast, MMP9 secretions increased 3.72- and 18.1-fold in HBs co-cultured with EPCs and EPCs, respectively. Immunofluorescence micrographs also showed that cMS with intercellular interactions between HBs and EPCs enhanced HGF and MMP9 expression (Figures 4D and S7). These results suggest that cMS elevate HGF secretion via intercellular communication between HBs and EPCs, whereas cMS elevates MMP9 secretion in EPCs without intercellular communication.

## Discussion

Our study investigated how cMS contributes to the expression and secretion of EMT-related proteins, including angiocrine signals, in HBs and EPCs using the OoC platform and hESCs. Since the previously reported OoC platform allows the application of cMS to cultured cells,^18^ we could apply cMS to differentiate hESCs. We confirmed that cMS did not alter the cellular population ratio of HBs to EPCs (8.5:1.5, estimated based on Figure S6A), which was similar to that of 15- to 24-week-old fetal human liver buds *in vivo* (HB: EC = 8.75:1.25).^27,41^ This suggests that the differentiation methods used for hESCs follow a hepatic developmental process.^16^

The obtained EPCs did not express CD31, a typical EC marker (Figures S1C and D), although another EC marker, KDR, was expressed (Figure 2C). Figures 2F and G show that KDR^+^ cells formed sprouting networks *in vitro* and expressed CD31. These results suggested that KDR^+^CD31^−^ cells could be EPCs and that CD31 was expressed only in differentiated endothelial cells. In contrast, previous studies suggested that KDR^+^CD31^−^ cells could be hepatic progenitor cells.^27,41^ KDR^+^CD31^−^ cells have both hepatic and endothelial progenitor cells. Therefore, combining KDR and CD31 is insufficient to distinguish between hepatic and endothelial progenitor cells to understand the origin of endothelial cells in the endoderm lineage. Thus, transcriptomic and proteomic analyses are required, followed by lineage tracing. In either case, endodermal cells, not the mesodermal cell lineage, could be a source of tissue-specific endothelial cells.

By inducing HBs and EPCs with cMS, we demonstrated that cMS enhanced EMT-related gene expression and HGF and MMP9 expression and secretion. GO analysis and key regulator analysis of upregulated DEG in D12_cMS^+^ by RNA-seq revealed that cMS upregulated genes related to tissue morphogenesis and vasculature development, and transcription factors STAT3 and CREB1, whose gene expression levels were not changed by cMS, and upregulated EMT-related genes, including HGF and MMP9.^42,43^ These transcription factors regulate the expression of EMT-related genes.^44,45^ Furthermore, the amounts of secreted HGF and MMP9 were enhanced by cMS, depending on the frequency and stretch. These results suggest that STAT3 and CREB1 are activated by calcium inflow^42,43,46^ as cMS causes continuous calcium ion inflow into the cytoplasm via the activation of mechanosensitive channels, such as PIEZO1 or TRPV4.^36,37^ Our gene expression profiles showed high gene expression levels for some PIEZO and TRP families, including TRPM4, TPRM7, and PIEZO1 (Figure S5). The activation of these channels may be related to the activation of STAT3 and CREB1, which contribute to the expression of EMT-related genes, including HGF and MMP9, in HBs and EPCs during vascular network formation. In addition, since the migration of HBs and EPCs during initial liver development is accompanied by EMT, cMS might contribute to vasculature development and liver bud formation.

By culturing HBs and EPCs separately, we demonstrated that intercellular communication between HBs and EPCs is necessary to enhance cMS-triggered angiocrine signals, such as HGF and MMP9 secretion. Although *HGF* expression in EPCs co-cultured with HBs and its secretion in EPCs alone were not enhanced by cMS, HGF expression and secretion were dramatically promoted by cMS when co-cultured with HBs. These results indicate that intercellular communication with HBs is involved in translation and secretion processes in EPCs. In contrast, both gene and protein expression of HGF in HBs was promoted by cMS when co-cultured with EPCs, suggesting that intercellular communication with EPCs is involved in HGF expression. In HBs, intercellular communication with EPCs may be involved in the upstream factor expression of STAT3 and CREB1 activated by cMS. cMS also promoted MMP9 secretion in EPCs co-cultured with HBs and EPCs alone but not in separately cultured HBs. The increased expression of MMP9 by cMS in separately cultured HBs was not larger than those in mixtures and mono-cultured EPCs; the intercellular communication with EPCs in HBs might also contribute to the activation of transcription factors of MMP9, such as STAT3 and CREB1. MMP9 secretion from the EPCs was also enhanced. Therefore, the intercellular communication of EPCs with HBs improves MMP9 secretion. Thus, enhanced expressions and secretions of these proteins by cMS through intercellular communication contribute to the vascular network of the liver lobule.

In this study, the PDMS membrane was ballooned and stretched to apply cMS to cultured cells. Stretching cells alters various cellular components and signaling pathways, including cytoskeletal remodeling, adherence junctions, focal adhesion formation, and extracellular protein rearrangement.^47^ PIEZO channels are major cellular mechanosensors associated with intracellular calcium regulation and contribute to developmental processes.^36^ It would be advantageous to perform live-cell imaging for calcium and lineage-tracing reporters under cMS to investigate the mechanisms of cMS in the developmental process. However, a limitation of this study is that our OoC platform was not suitable for observing cells using faster fluorescent microscopes (a light-sheet microscope or a spinning-disk confocal microscope) owing to the multiple PDMS layers required to create a long distance from the samples to an objective lens. Therefore, next-generation OoC platforms must fulfill the requirements of microscopy for observing cells to elucidate the underlying mechanisms by which cMS contributes to liver developmental processes.

In addition to OoC platforms, current *in vitro* hepatic differentiation processes should be considered to understand the contribution of cMS. Although HBs and EPCs were simultaneously differentiated from hESCs by mimicking the embryonic hepatic developmental process in this study, we also need to consider the intercellular communication with other cells or tissues, such as the mesenchyme, to completely understand the formation of a vascular network in the liver lobule. Many research groups, including ours, have supplemented BMP4 and FGF during hepatic differentiation because BMP4 and FGF are secreted from the septum transversum mesenchyme and cardiac mesoderm, respectively.^48–50^ Thus, a multicell culturing system would be beneficial for studying the embryonic tissue developmental process, and OoC could be utilized to manage mechanical stimuli on organoids.

In conclusion, cMS activates angiocrine signals in HBs and EPCs during HB commitment via intercellular communication between HBs and EPCs. Our approach and results provide insight into the mechanisms of liver bud formation and tissue engineering to obtain fully functional liver tissues with a vascular network.

## Acknowledgments

We thank Ms. Shiho Terada for assisting with the hESCs culture, Ms. Kyoko Sawada for arranging tables of antibodies used, Kyoto University Graduate School of Biostudies Laboratory for Biological Dynamics, and Mr. Daiya Ohara for supporting the RNA-seq and analysis.

We thank 9^th^ World Congress of Biomechanics 2022 Taipei for the presentation of preliminary data.

## Supporting information

Additional Supporting Information is available online at the end of this article.

## Data availability

RNA-seq data were deposited in the NCBI Gene Expression Omnibus under Accession Number: GSE232118.

## Abbreviations

cMS: cyclic mechanical stretching
CYP3A: cytochrome P450 3A
D9: day 9
D12: day 12
DEG: differentially expressed gene
EC: endothelial cell
EMT: epithelial-to-mesenchymal transition
EPC: endothelial progenitor cells
FACS: fluorescence-activated cell sorter
FOXA2: forkhead box A2
HB: hepatoblast
HE: hepatic endoderm
hESCs: human embryonic pluripotent stem cells
HHEX: hematopoietically-expressed homeobox protein
hiPSCs: human induced pluripotent stem cells
HNF4α: hepatocyte nuclear factor 4α
hPSCs: human pluripotent stem cells
KRT19: keratin 19
MMP9: matrix metallopeptidase 9
NEAA: MEM Non-essential Amino Acid Solution without L-glutamine, liquid, sterile-filtered Bioreagent suitable for cell-culture
OoC: organ-on-a-chip
PDMS: polydimethylsiloxane
RNA-seq: RNA sequencing
RPMI: Roswell Park Memorial Institute
SD: standard deviation
SOX17: SRY-box transcription factor 17
TPM: transcripts per kilobase million
TRPV: transient receptor potential vanilloid

## References

1. Prior N, Inacio P, Huch M. Liver organoids: from basic research to therapeutic applications. Gut. 2019:68(12):2228–2237.

2. Ramli MN Bin, Lim YS, Koe CT, et al. Human Pluripotent Stem Cell-Derived Organoids as Models of Liver Disease. Gastroenterology. 2020;159(4):1471–1486.e12.

3. Sun XC, Kong DF, Zhao J, Faber KN, Xia Q, He K. Liver organoids: established tools for disease modeling and drug development. Hepatol Commun. 2023;7(4):e0105.

4. Thomson JA. Embryonic Stem Cell Lines Derived from Human Blastocysts. Science 1998;282(5391):1145–1147.

5. Takahashi K, Tanabe K, Ohnuki M, et al. Induction of Pluripotent Stem Cells from Adult Human Fibroblasts by Defined Factors. Cell. 2007;131(5):861–872.

6. Wesley BT, Ross ADB, Muraro D, et al. Single-cell atlas of human liver development reveals pathways directing hepatic cell fates. Nat Cell Biol. 2022;24(10):1487–1498.

7. Ouchi R, Koike H. Modeling human liver organ development and diseases with pluripotent stem cell-derived organoids. Front Cell Dev Biol. 2023;11:1133534.

8. Hofer M, Lutolf MP. Engineering organoids. Nat Rev Mater. 2021;6(5):402–420.

9. Takebe T, Sekine K, Kimura M, et al. Massive and Reproducible Production of Liver Buds Entirely from Human Pluripotent Stem Cells. Cell Rep. 2017;21(10):2661–2670.

10. Mun SJ, Ryu JS, Lee MO, et al. Generation of expandable human pluripotent stem cell-derived hepatocyte-like liver organoids. J Hepatol. 2019;71(5):970–985.

11. Si-Tayeb K, Lemaigre FP, Duncan SA. Organogenesis and Development of the Liver.Dev Cell. 2010;18(2):175–189.

12. Gruppuso PA, Sanders JA. Regulation of liver development: Implications for liver biology across the lifespan. J Mol Endocrinol. 2016;56(3):R115–R125.

13. Matsumoto K, Yoshitomi H, Rossant J, Zaret KS. Liver organogenesis promoted by endothelial cells prior to vascular function. Science. 2001;294(5542):559–563.

14. Gordillo M, Evans T, Gouon-Evans V. Orchestrating liver development. Development. 2015;142(12):2094–2108.

15. Lorenz L, Axnick J, Buschmann T, et al. Mechanosensing by β1 integrin induces angiocrine signals for liver growth and survival. Nature. 2018;562(7725):128–132.

16. Kamei K ichiro, Yoshioka M, Terada S, Tokunaga Y, Chen Y. Three-dimensional cultured liver-on-a-Chip with mature hepatocyte-like cells derived from human pluripotent stem cells. Biomed Microdevices. 2019;21(3):73.

17. Huh D, Matthews BD, Mammoto A, Montoya-Zavala M, Hsin HY, Ingber DE. Reconstituting Organ-Level Lung Functions on a Chip. Science. 2010;328(5986):1662–1668.

18. Yoshimoto K, Minier N, Yang J, et al. Recapitulation of Human Embryonic Heartbeat to Promote Differentiation of Hepatic Endoderm to Hepatoblasts. Front Bioeng Biotechnol. 2020;8:568092.

19. Kamei K ichiro, Mashimo Y, Koyama Y, et al. 3D printing of soft lithography mold for rapid production of polydimethylsiloxane-based microfluidic devices for cell stimulation with concentration gradients. Biomed Microdev. 2015;17(2):36.

20. Kim D, Langmead B, Salzberg SL. HISAT: A fast spliced aligner with low memory requirements. Nat Methods. 2015;12(4):357–360.

21. Liao Y, Smyth GK, Shi W. FeatureCounts: An efficient general purpose program for assigning sequence reads to genomic features. Bioinformatics. 2014;30(7):923–930.

22. Ge SX, Son EW, Yao R. iDEP: An integrated web application for differential expression and pathway analysis of RNA-Seq data. BMC Bioinformatics. 2018;19(1):1–24.

23. Zhou Y, Zhou B, Pache L, et al. Metascape provides a biologist-oriented resource for the analysis of systems-level datasets. Nat Commun. 2019;10(1):1523.

24. Han H, Cho JW, Lee S, et al. TRRUST v2: An expanded reference database of human and mouse transcriptional regulatory interactions. Nucleic Acids Res. 2018;46(D1):D380–D386.

25. Carpenter AE, Jones TR, Lamprecht MR, et al. CellProfiler: Image analysis software for identifying and quantifying cell phenotypes. Genome Biol. 2006;7(10):R100.

26. Otsu N. A Threshold Selection Method from Gray-Level Histograms. IEEE Trans Syst Man Cybern. 1979;9(1):62–66.

27. Goldman O, Han S, Hamou W, et al. Endoderm generates endothelial cells during liver development. Stem Cell Reports. 2014;3(4):556–565.

28. Hannan NRF, Segeritz CP, Touboul T, Vallier L. Production of hepatocyte-like cells from human pluripotent stem cells. Nat Protoc. 2013;8(2):430–437.

29. Seto Y, Inaba R, Okuyama T, Sassa F, Suzuki H, Fukuda J. Engineering of capillary-like structures in tissue constructs by electrochemical detachment of cells. Biomaterials. 2010;31(8):2209–2215.

30. Farrell J, Kelly C, Rauch J, et al. HGF induces epithelial-to-mesenchymal transition by modulating the mammalian Hippo/MST2 and ISG15 pathways. J Proteome Res. 2014;13(6):2874–2886.

31. Sullivan NJ, Sasser AK, Axel AE, et al. Interleukin-6 induces an epithelialmesenchymal transition phenotype in human breast cancer cells. Oncogene. 2009;28(33):2940–2947.

32. Fang R, Zhang G, Guo Q, et al. Nodal promotes aggressive phenotype via Snail-mediated epithelial-mesenchymal transition in murine melanoma. Cancer Lett. 2013;333(1):66–75.

33. Lindsley RC, Gill JG, Murphy TL, et al. Mesp1 coordinately regulates cardiovascular fate restriction and epithelial-mesenchymal transition in differentiating ESCs. Cell Stem Cell. 2008;3(1):55–68.

34. Shirakihara T, Horiguchi K, Miyazawa K, et al. TGF-β regulates isoform switching of FGF receptors and epithelial-mesenchymal transition. EMBO J. 2011;30(4):783–795.

35. Bhatia S, Wang P, Toh A, Thompson EW. New Insights Into the Role of Phenotypic Plasticity and EMT in Driving Cancer Progression. Front Mol Biosci. 2020;7:71.

36. He L, Si G, Huang J, Samuel ADT, Perrimon N. Mechanical regulation of stem-cell differentiation by the stretch-activated Piezo channel. Nature. 2018;555(7694):103–106.

37. Zhao Q, Zhou H, Chi S, et al. Structure and mechanogating mechanism of the Piezo1 channel. Nature. 2018;554(7693):487–492.

38. Suárez-Causado A, Caballero-Díaz D, Bertrán E, et al. HGF/c-Met signaling promotes liver progenitor cell migration and invasion by an epithelial-mesenchymal transition-independent, phosphatidyl inositol-3 kinase-dependent pathway in an in vitro model. Biochim Biophys Acta Mol Cell Res. 2015;1853(10):2453–2463.

39. Quondamatteo F, Knittel T, Mehde M, Ramadori G, Herken R. Matrix metalloproteinases in early human liver development. Histochem Cell Biol. 1999;112(4):277–282.

40. Margagliotti S, Clotman F, Pierreux CE, et al. Role of metalloproteinases at the onset of liver development. Dev Growth Differ. 2008;50(5):331–338.

41. Goldman O, Han S, Sourrisseau M, et al. KDR identifies a conserved human and murine hepatic progenitor and instructs early liver development. Cell Stem Cell. 2013;12(6):748–760.

42. Miras-Portugal MT, Gomez-Villafuertes R, Gualix J, et al. Nucleotides in neuroregeneration and neuroprotection. Neuropharmacology. 2016;104:243–254.

43. Sheng M, Thompson MA, Greenberg ME. CREB: A Ca2+-regulated transcription factor phosphorylated by calmodulin-dependent kinases. Science. 1991;252(5011):1427–1430.

44. Du P, Zeng H, Xiao Y, et al. Chronic stress promotes EMT-mediated metastasis through activation of STAT3 signaling pathway by miR-337-3p in breast cancer. Cell Death Dis. 2020;11(9):761.

45. Won SY, Park JJ, Shin EY, Kim EG. PAK4 signaling in health and disease: defining the PAK4–CREB axis. Exp Mol Med. 2019;51(2):1–9.

46. Murase S, Mckay RD. Neuronal activity-dependent STAT3 localization to nucleus is dependent on Tyr-705 and Ser-727 phosphorylation in rat hippocampal neurons. Eur J Neurosci. 2014;39(4):557–565.

47. Przybyla L, Muncie JM, Weaver VM. Mechanical Control of Epithelial-to-Mesenchymal Transitions in Development and Cancer. Annu Rev Cell Dev Biol. 2016;32:527–554.

48. Wang J, Rhee S, Palaria A, Tremblay KD. FGF signaling is required for anterior but not posterior specification of the murine liver bud. Dev Dyn. 2015;244(3):431–443.

49. Tremblay KD, Zaret KS. Distinct populations of endoderm cells converge to generate the embryonic liver bud and ventral foregut tissues. Dev Biol. 2005;280(1):87–99.

50. Palaria A, Angelo JR, Guertin TM, Mager J, Tremblay KD. Patterning of the hepato-pancreatobiliary boundary by BMP reveals heterogeneity within the murine liver bud. Hepatology. 2018;68(1):274–288.

