## Supplementary information for "Cyclic mechanical stretching stimuli promotes angiocrine signals during *in vitro* liver bud formation from human pluripotent stem cells"

**Supplementary Figures**

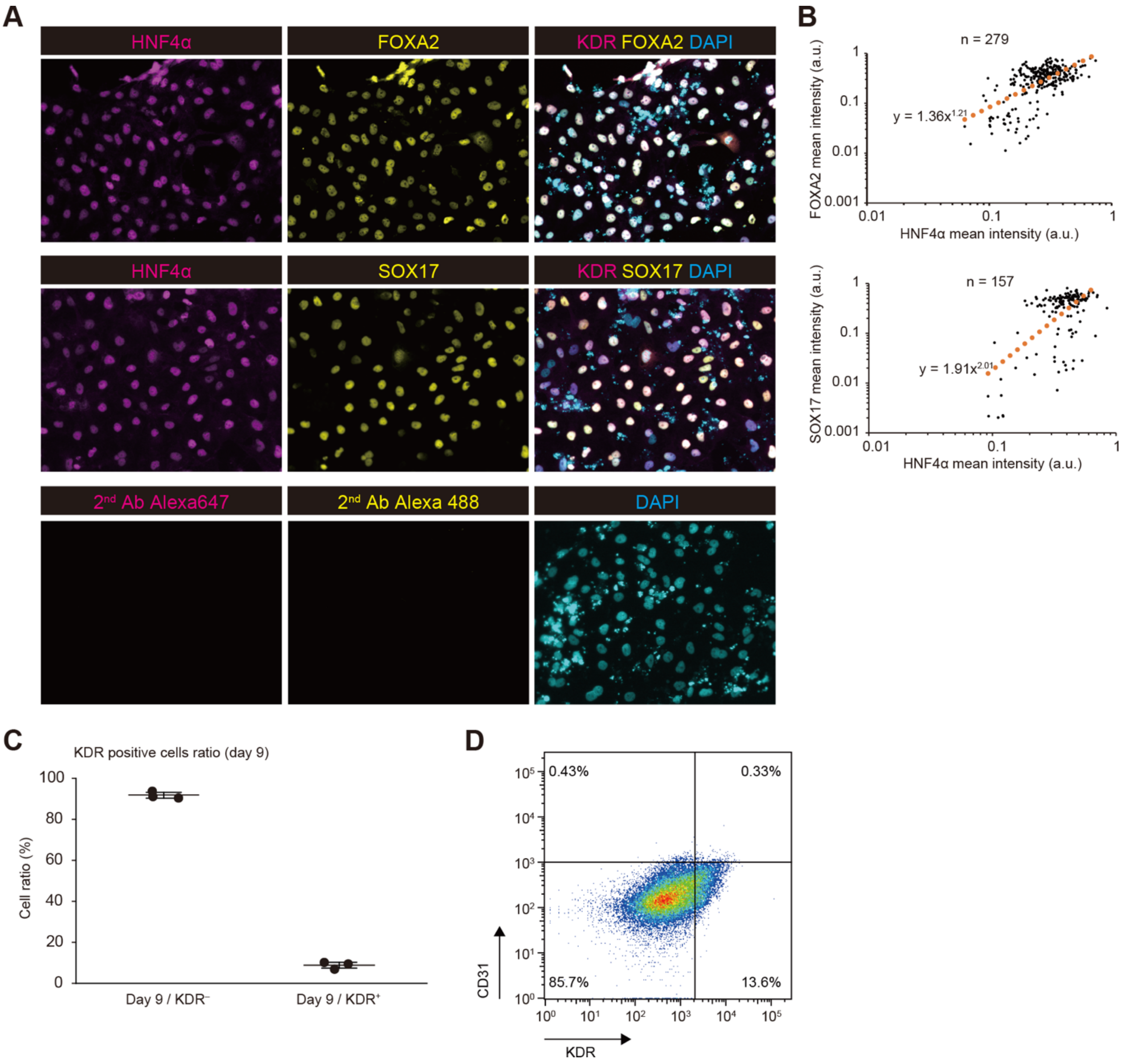


**Figure S1. Characterization of the co-cultured HEs and EPCs derived from hPSCs at D9.** (A) Immunofluorescence micrographs of D9 cells of the mixture stained with the HE markers (FOXA2, yellow; SOX17, yellow; HNF4α, magenta) and the nuclei (DAPI, cyan) and their negative controls. Scale bars represent 100 µm. (B) Quantitative single-cell profiling of each protein at D9 based on the result of the immunofluorescence staining. The approximate lines were fitted to quantitative single-cell profiled results to show the correlative relationships. (C) The KDR-positive and -negative cells ratio with flow cytometry at D9. (D) Flow cytometry analysis of a mixture with an anti-KDR antibody and an anti-CD31 antibody at D9.

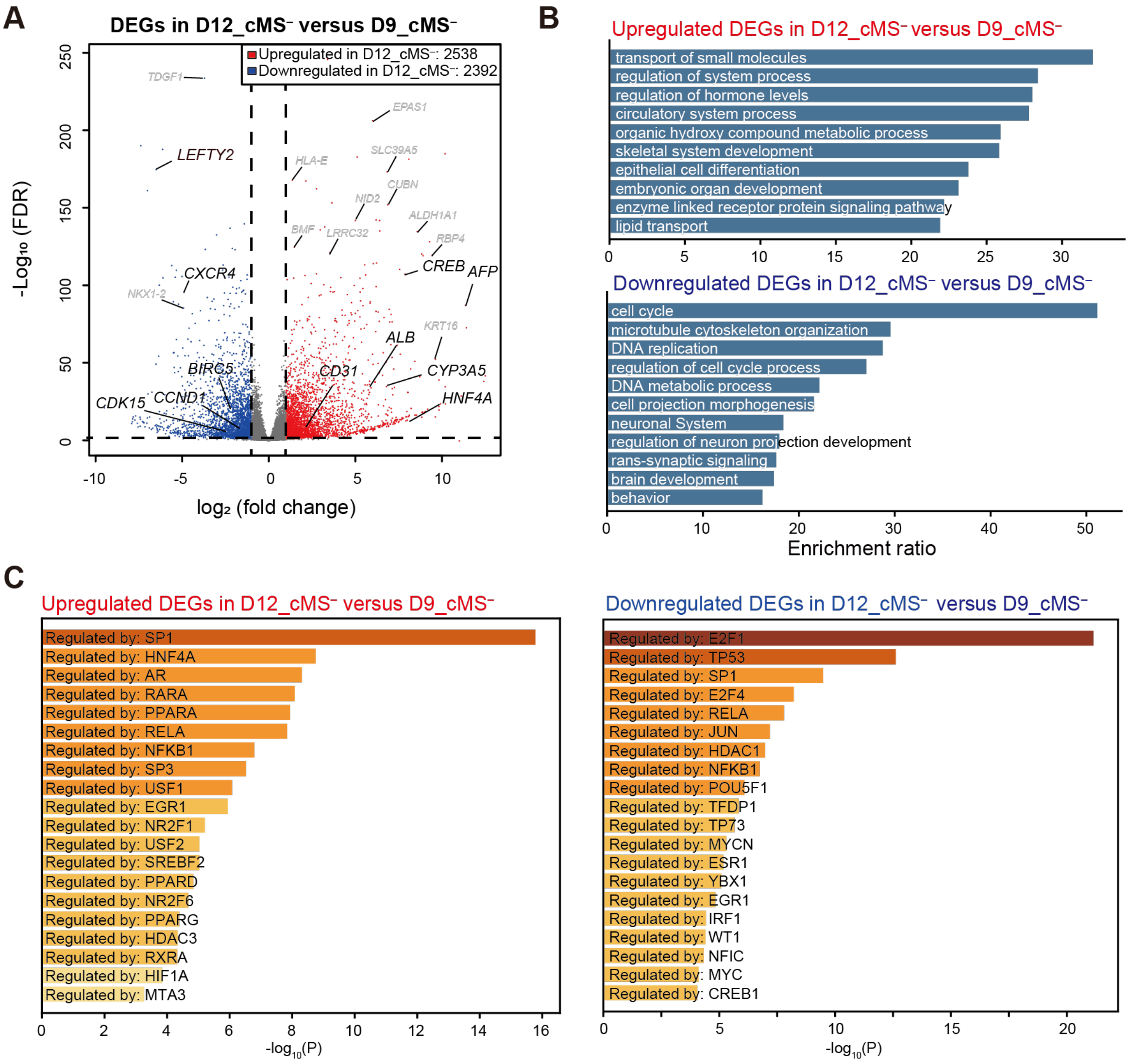


**Figure S2. Comparative transcriptional profiling of hESC-derived cells at D9_cMS$^{-}$ and D12 without cMS (D12_cMS$^{-}$).** (A) A volcano plot of RNA-seq by comparing D9_cMS$^{-}$ and D12_cMS$^{-}$. (B) Gene ontology terms based on 2538 upregulated and 2392 downregulated DEGs in D12_cMS$^{-}$ compared to those in D9_cMS$^{-}$. The upregulated DEGs included HB and EC markers, such as *CYP3A5*, *HNF4*, *AFP, ALB*, and *CD31*, while the downregulated genes contained endoderm markers (*CXCR4*) and cell division-related genes (*CDK15*, *CCND1*, and *BIRC*). (C) Identification of transcription factors regulating DEGs.

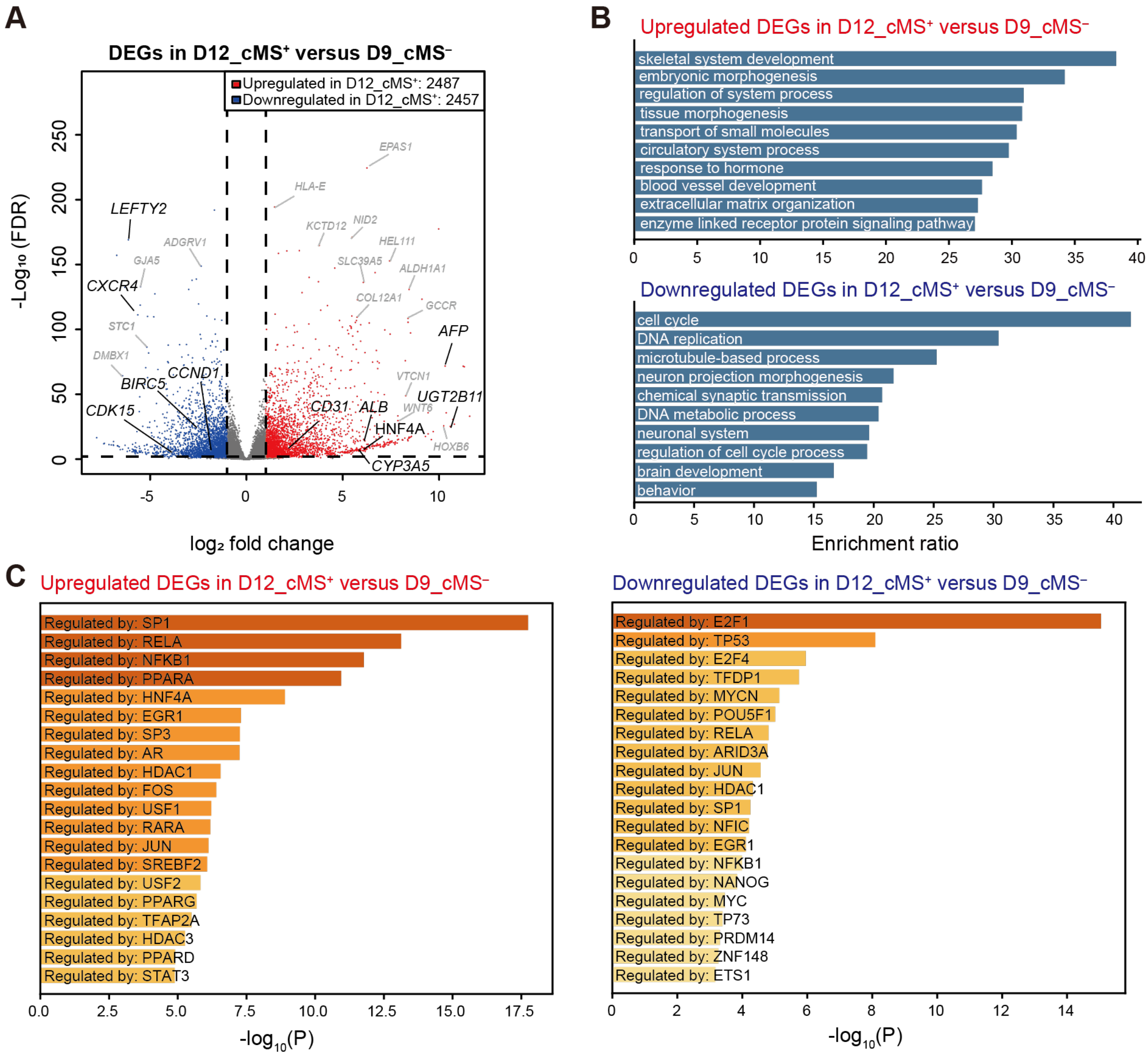


**Figure S3. Comparative transcriptional profiling of cells derived from hESCs at D9_cMS$^{-}$ and D12 with cMS (D12_cMS$^{+}$).** (A) A volcano plot of RNA-seq by comparing D9_cMS$^{-}$ and D12_cMS$^{+}$. (B) Gene ontology terms based on 2487 upregulated and 2457 downregulated DEGs in D12_cMS$^{+}$ compared to those in D9_cMS$^{-}$. The upregulated DEGs in D12_cMS$^{+}$ had HB and EC markers such as *CYP3A5*, *HNF4*, *AFP, ALB*, and *CD31*. The downregulated DEGs in D12_cMS$^{+}$ had endoderm markers such as *CXCR4* and cell division-related genes such as *CDK15*, *CCND1*, and *BIRC*. (C) Identification of transcription factors regulating DEGs.

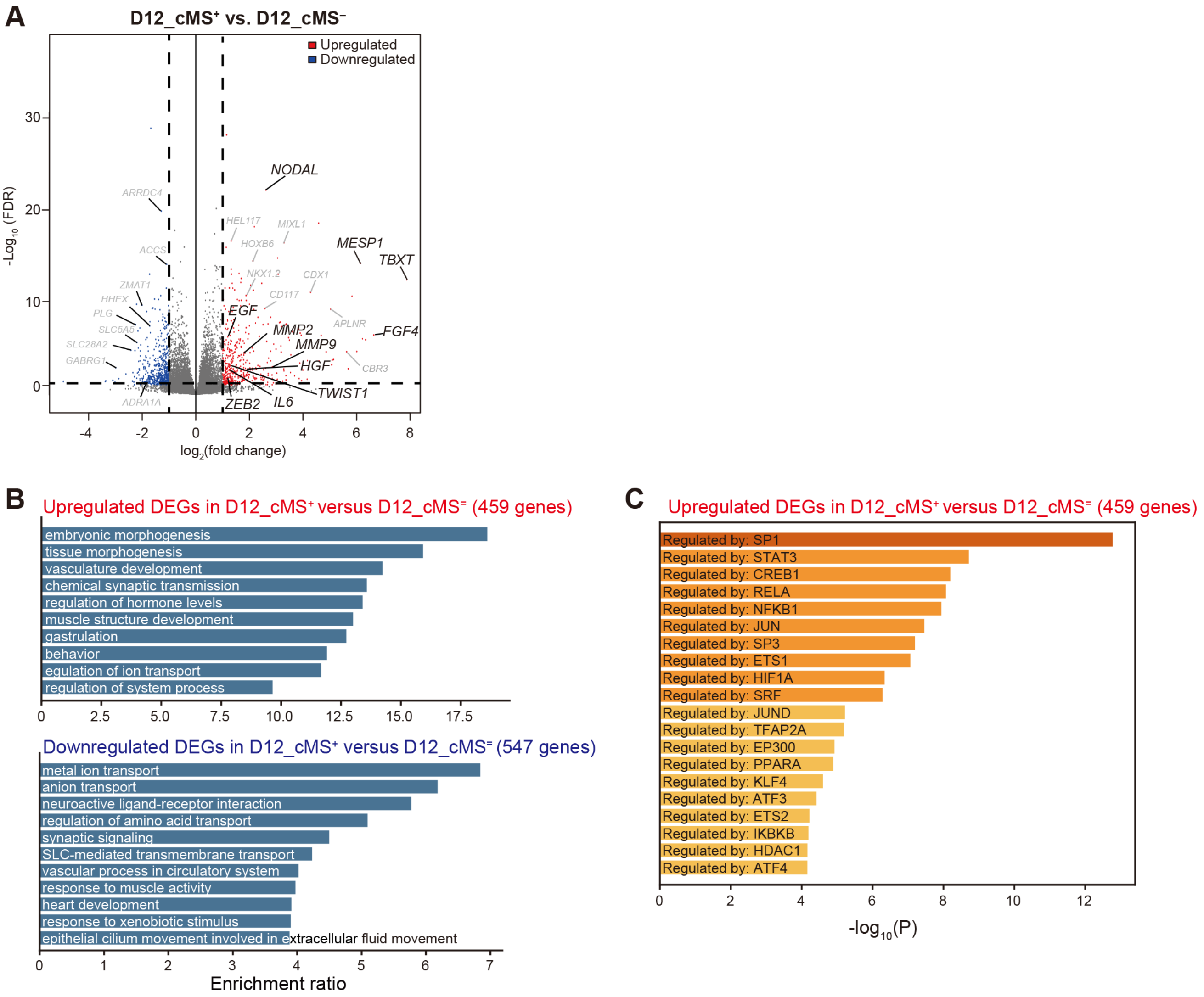


**Figure S4. Comparative transcriptional profiling of cells derived from hESCs at D12 with and without cMS (D12_cMS$^{+}$ and D12_cMS$^{-}$, respectively).** (A) A volcano plot of transcriptional profiling results with DEGs from the comparison of D12_cMS$^{-}$ and D12_cMS$^{+}$. Red and blue plots show DEGs identified with false discovery rate (FDR) < 0.05 and |$\log_2$ fold change (FC)| > 1. (B) Gene ontology terms based on 384 upregulated and 378 downregulated DEGs in D12_cMS$^{+}$ compared to those in D12_cMS$^{-}$. We found that the upregulated DEGs in D12_cMS$^{+}$ have a collection of the EMT-related genes, including angiocrine-related genes (*HGF*, *MMP2*, *NODAL*, *MMP9*, *MESP1*, *IL6*, *FGF4*, *ZEB2* and *TWIST1*) (1–6), while the downregulated DEGs contained endoderm markers (*CXCR4*) and ion transport-related genes (*SLC5A5*, *SLC6A3*, and *SLC9A2*). (C) Identification of transcription factors regulating DEGs.

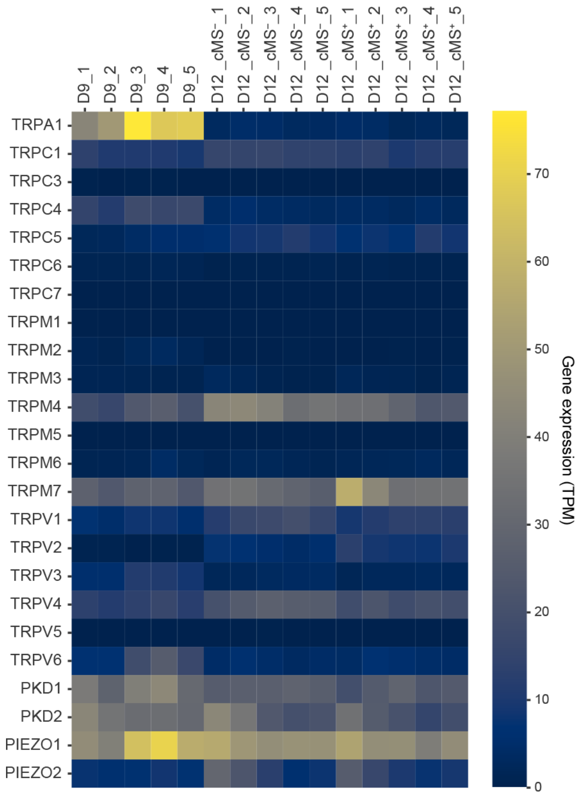


**Figure S5. Heatmap of mechanosensitive TRP and PIEZO families of D9_cMS$^-$, D12_cMS$^-$, and D12_cMS$^+$.**

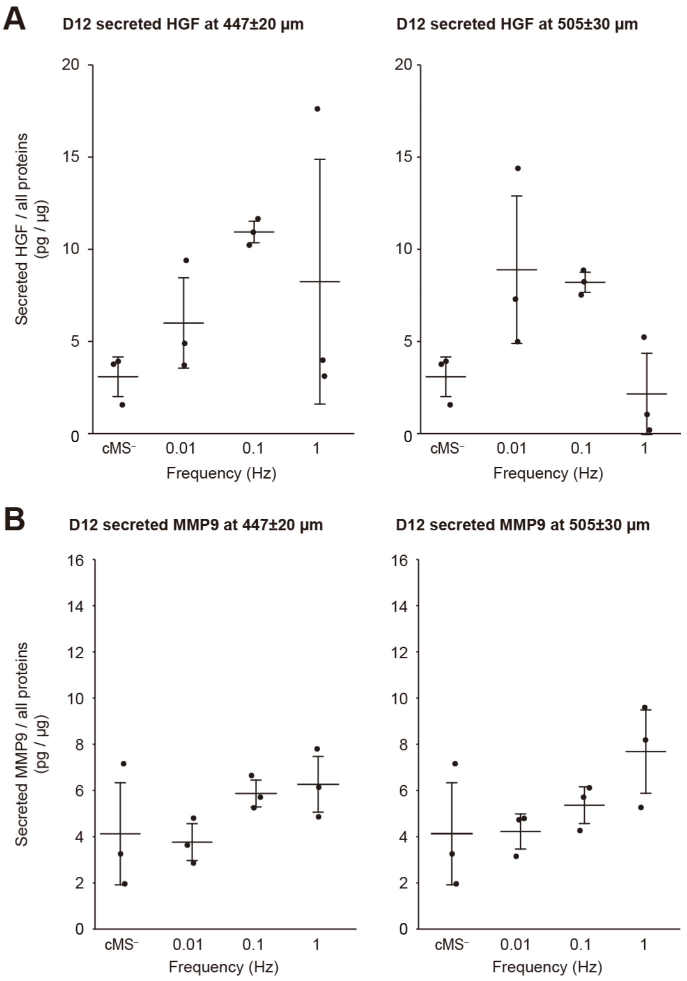


**Figure S6. Amounts of secreted HGF and MMP9 in the mixtures with 0.01, 0.1, and 1 Hz cMS and without cMS determined by ELISA.** (A) Amounts of secreted HGF in the mixtures with 0.01, 0.1, and 1 Hz cMS at 447 ± 20 µm and 505 ± 30 µm of the membrane displacement and in the mixture without cMS. (B) Amounts of secreted MMP9 in the mixtures with 0.01, 0.1, and 1 Hz cMS at 447 ± 20 µm and 505 ± 30 µm of the membrane displacement and in the mixture without cMS. All data represent mean ± SD.

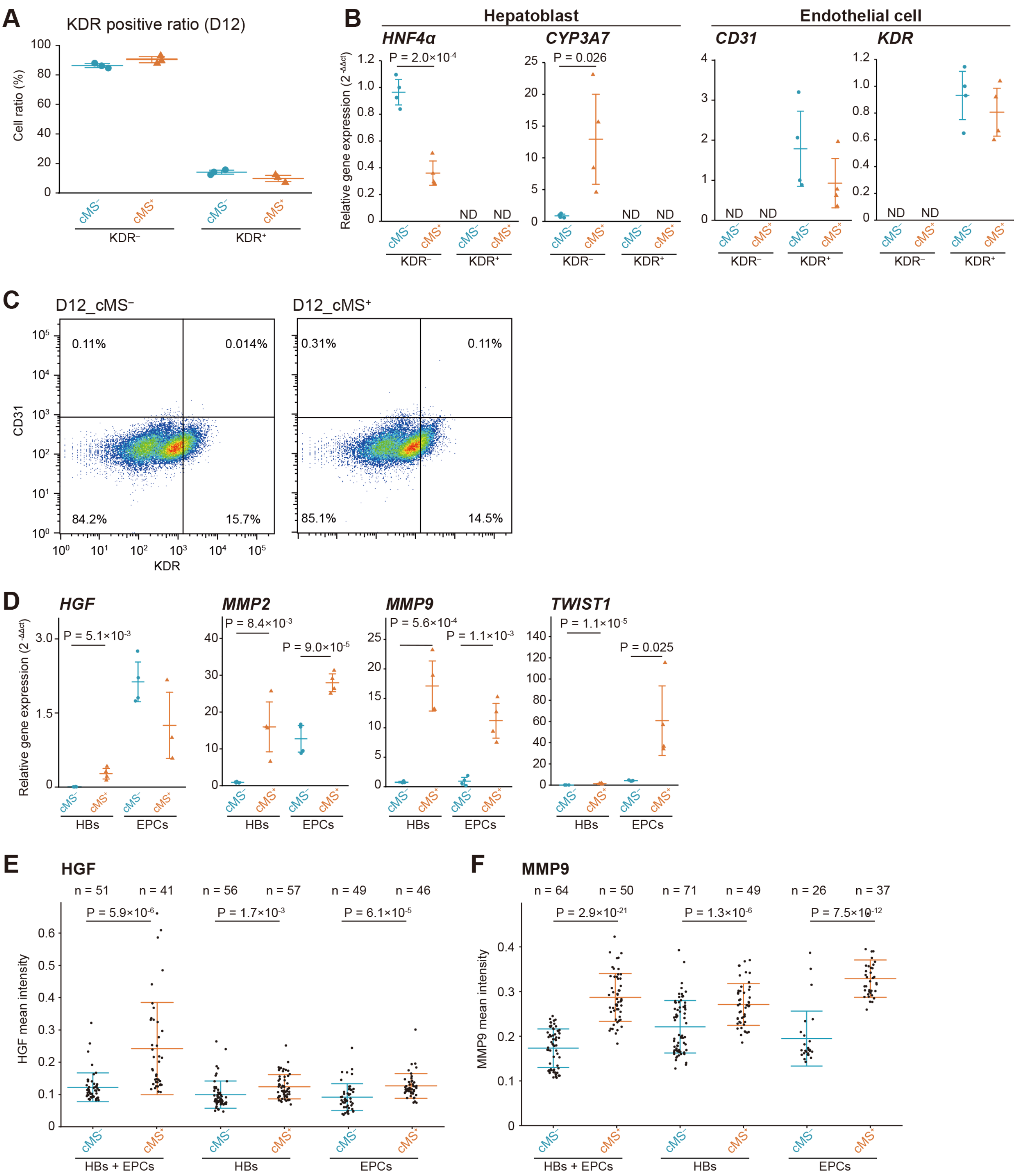


**Figure S7. Characterization of the co-cultured HBs and EPCs with and without cMS at D12.** (A) The percentile of KDR$^{-}$ cells and KDR$^{+}$ cells determined by flow cytometry in D12_cMS$^{-}$ and D12_cMS$^{+}$. (B) RT-qPCR to determine the expression of HB (*HNF4a* and *CYP3A7*) and EC (*CD31* and *KDR*) markers in HBs and EPCs by applying cMS. For HB markers, only KDR$^{-}$ cells expressed *CYP3A7* and *HNF4A*. By applying cMS, *CYP3A7* expression in KDR$^{-}$ cells was increased by 14.7-folds, while *HNF4A* expression in KDR$^{-}$ cells was reduced by 2.68-folds. In contrast, for EC markers, only KDR$^{+}$cells expressed *KDR* and *CD31*, and cMS did not alter the expression of *KDR* and *CD31* in both $KDR^{-}$ and $KDR^{+}$cells Data represent means ± SD. The significance value was calculated using a two-tailed unpaired Student's *t*-test (n=4). (C) Flow cytometry analysis of $D12\_cMS^{-}$ and $D12\_cMS^{+}$ with HB (KDR) and EC (CD31) markers. (D) RT-qPCR to determine the expression of EMT-associated genes (*HGF*, *MMP9*, *MMP2*, and *TWIST1*) in HBs and EPCs by applying cMS. *MMP2* also showed a similar trend with MMP9, and *MMP2* expression in $HBs\_cMS^{+}$ and $EPCs\_cMS^{+}$ showed 17.71- and 2.20-fold increases, respectively, with cMS. For *TWIST1*, $HBs\_cMS^{+}$ and $EPCs\_cMS^{+}$ expression showed 12.51- and 14.50-fold increases with cMS but not in HBs. Data represent means ± SD. The significance value was calculated using a two-tailed unpaired Student's *t*-test (n=4). (E, F) Quantitative single-cell profiling of HGF and MMP9 expression in the co-cultured and mono-cultured HBs and EPCs based on the micrographs shown in Figure 4D. Data represent means ± SD. The significance value was calculated using Welch's test. By applying cMS, the expression of HGF was 1.98-fold increased in the co-cultured cells, although HGF expressions in HBs and EPCs were 1.38- and 1.24-fold increased. The increases in the average expression levels of MMP9 by cMS in co-cultured and mono-cultured EPCs (1.69- and 1.65-fold) were higher than that in mono-cultured HBs (1.22-fold).

**Table S1. DEGs of D12_cMS$^{-}$ compared with D9_cMS$^{-}$**

**Table S2. DEGs of D12_cMS$^{+}$ compared with D9_cMS$^{-}$**

**Table S3. DEGs of D12_cMS$^{+}$ compared with D12_cMS$^{-}$**

**Table S4. DEGs in D12_cMS$^{+}$ but not in D12_cMS nor D9_cMS$^{-}$**

**Table S5. Antibodies used in flow cytometry**

| Name | Company | Catalog number | Dilution |
|---|---|---|---|
| Alexa Fluor 647 Mouse Anti-Human CD309 (VEGFR-2) | Biosciences | 560495 | 1:50 |
| Alexa Fluor 488 anti-human CD31 | BioLegend | 303110 | 1:50 |
| Alexa 647 Mouse IgG1κ isotype control | BD Bioscience | 557783 | |
| Alexa Fluor 488 mouse IgG1κ isotype control | BD Bioscience | 557782 | 1:50 |

**Table S6. The primers used in the real-time RT-PCR analysis**

| Gene Symbol | Primers (forward/reverse; 5’ to 3’) |
|---|---|
| *FOXA2* | GGAACACCACTACGCCTTCAAC/AGTGCATCACCTGTTCGTAGGC |
| *CD31* | AAGTGGAGTCCAGCCGCATATC/ATGGAGCAGGACAGGTTCAGTC |
| *KDR* | GGAACCTCACTATCCGCAGAGT/CCAAGTTCGTCTTTTCCTGGGC |
| *TWIST1* | GCCAGGTACATCGACTTCCTCT/TCCATCCTCCAGACCGAGAAGG |
| *HGF* | GAGAGTTGGGTTCTTACTGCACG/CTCATCTCCTCTTCCGTGGACA |
| *GAPDH* | GCACCGTCAAGGCTGAGAAC/TGGTGAAGACGCCAGTGGA |
| *CYP3A7* | AAGTCTGGGGTATTTATGACT/CGCTGGTGAATGTTGGAGAC |
| *HNF4A* | GAGCTGCAGATCGATGACAA/TACTGGCGGTCGTTGATGTA |
| *HHEX* | CCAGGTGAGATTCTCCAACGAC/CTCCATTTAGCGCGTCGATTCTG |
| *KRT19* | AGCTAGAGGTGAAGATCCGCGA/GCAGGACAATCCTGGAGTTCTC |
| *MMP2* | TACAGGATCATTGGCTACACACC/GGTCACATCGCTCCAGACT |
| *SOX17* | AGGAAATCCTCAGACTCCTGGGTT/CCCAAACTGTTCAAGTGGCAGACA |

**Table S7. The antibodies used in immunocytochemistry**

| Name | Company | Catalog number | Dilution |
|---|---|---|---|
| MMP9 Antibody | Bioss | bs-0397R | 1:500 |
| HHEX Antibody | R&D systems | MAB83771-100 | 1:200 |
| SOX17 Antibody | R&D systems | MAB1924 | 1:200 |
| HGF Antibody | Protein tech | 26881-1-AP | 1:500 |
| KRT19 Antibody | Thermo Fisher | 61-7300 | 1:500 |
| FOXA2 Antibody | R&D systems | AF2400 | 1:200 |
| KDR Antibody | Abcam | ab2349 | 1:500 |
| HNF4α Antibody | Cell signaling | mAb #3113 | 1:200 |
| CD31 Antibody | R&D systems | AF3628 | 1:500 |
| AlexaFluor 488 donkey anti-mouse IgG (H + L) | Jackson ImmunoResearch | 715-545-150 | 1:1000 |
| AlexaFluor 647 donkey anti-rabbit IgG (H + L) | Jackson ImmunoResearch | 711-606-152 | 1:1000 |
| AlexaFluor 488 donkey Anti-goat IgG | Abcam | Ab150129 | 1:1000 |

**Supplementary References**


1. Farrell J, Kelly C, Rauch J, Kida K, García-Muñoz A, Monsefi N, et al. HGF induces epithelial-to-mesenchymal transition by modulating the mammalian Hippo/MST2 and ISG15 pathways. J Proteome Res. 2014;13:2874–2886.
2. Sullivan NJ, Sasser AK, Axel AE, Vesuna F, Raman V, Ramirez N, et al. Interleukin-6 induces an epithelial-mesenchymal transition phenotype in human breast cancer cells. Oncogene. 2009;28:2940–2947.
3. Fang R, Zhang G, Guo Q, Ning F, Wang H, Cai S, et al. Nodal promotes aggressive phenotype via Snail-mediated epithelial-mesenchymal transition in murine melanoma. Cancer Lett. 2013;333:66–75.
4. Lindsley RC, Gill JG, Murphy TL, Langer EM, Cai M, Mashayekhi M, et al. Mesp1 coordinately regulates cardiovascular fate restriction and epithelial-mesenchymal transition in differentiating ESCs. Cell Stem Cell. 2008;3:55–68.
5. Shirakihara T, Horiguchi K, Miyazawa K, Ehata S, Shibata T, Morita I, et al. TGF-β regulates isoform switching of FGF receptors and epithelial-mesenchymal transition. EMBO J. 2011;30:783–795.
6. Bhatia S, Wang P, Toh A, Thompson EW. New Insights Into the Role of Phenotypic Plasticity and EMT in Driving Cancer Progression. Front Mol Biosci. 2020;7:1–18.
